# Improved prediction of behavioral and neural similarity spaces using pruned DNNs

**DOI:** 10.1101/2021.07.08.451521

**Authors:** Homa Priya Tarigopula, Scott Laurence Fairhall, Uri Hasson

## Abstract

Deep Neural Networks (DNNs) have become an important tool for modeling brain and behaviour. One key area of interest has been to apply these networks to model human similarity judgements. Several previous works have used the embeddings from the penultimate layer of vision DNNs and showed that a reweighting of these features improves the fit between human similarity judgments and DNNs. These studies underline the idea that these embeddings form a good basis set but lack the correct level of salience. Here we re-examined the grounds for this idea and on the contrary, we hypothesized that these embeddings, beyond forming a good basis set, also have the correct level of salience to account for similarity judgments. It is just that the huge dimensional embedding needs to be pruned to select those features relevant for the considered domain for which a similarity space is modeled. In Study 1 we supervised DNN pruning based on a subset of human similarity judgments. We found that pruning: *i*) improved out-of-sample prediction of human similarity judgments from DNN embeddings, *ii*) produced better alignment with WordNet hierarchy, and *iii*) retained much higher classification accuracy than reweighting. Study 2 showed that pruning by neurobiological data is highly effective in improving out-of-sample prediction of brain-derived representational dissimilarity matrices from DNN embeddings, at times fleshing out isomorphisms not otherwise observable. Pruning supervised by human brain/behavior therefore effectively identifies alignable dimensions of semantic knowledge between DNNs and humans and constitutes an effective method for understanding the organization of knowledge in neural networks.

## 1 Introduction

### 1.1 State of the art

Deep Neural Networks for computer vision are now routinely used as predictive models of human brain and behavior (e.g., Cichy and Kaiser, 2019). A key question is whether the networks develop representations organized according to latent dimensions similar to those that structure human knowledge. For a given set of images this issue can be addressed by *i*) soliciting pairwise similarity judgments from humans; *ii*) computing cosine similarity for the same set from DNN embeddings, and *iii*) evaluating the correspondence between the two similarity matrices. The magnitude of this correspondence is reported as an *R*^2^ value and frequently referred to as second-order-isomorphism (*2OI*, Kriegeskorte et al., 2008). Practically, it reflects the network’s capacity to predict human similarity judgments. There is no need to use explicit similarity judgements; these can be replaced by any procedure that outputs a similarity space from human behavior or neural activity.

There is substantial heterogeneity in 2OI *R*^2^ values reported when relating DNN embeddings to human similarity judgments. Peterson et al. (2018) analyzed embeddings from the penultimate layer of VGG-19 (Simonyan and Zisserman, 2014) and reported 2OI for six image-sets, each set drawn from a different semantic category such as ANIMALS or FRUITS (a VARIOUS combined across categories). Reported values were in the range of *R*^2^ = 0.2 − 0.6. King et al. (2019) reported similar values for object and scene categories with a maximal Spearman’s Rho value of 0.56 (*R*^2^ not reported). Groen et al. (2018), studying more complex scenes, reported 2OI values of *R*^2^ = 0.07, (*r* = 0.26). A large scale analysis based on estimating psychological embeddings from 50,000 stimuli (Roads and Love, 2021) reported Spearman Rho values between 0.02 and 0.36, across 12 DNN architectures.

However, Peterson et al. were able to increase 2OI’s *R*^2^ by reweighting the activation (output) of each node in a DNN’s penultimate layer (without changing weights or biases), a process that can be considered as modifying each feature’s salience. This was achieved by maximizing the fit between the human and DNN’s similarity matrices through linear regression. Practically, the fit between human similarity judgment (*s*) and DNN-pairwise-similarity for any two images *k*, *j* was defined as 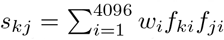. Here, 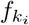 and 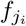 are the the values of feature *i* (of *n* = 4096) for image *k* and image *j*. The estimated pair-wise similarity *s*_*kj*_ then is just the sum of these products across all *n* = 4096 features. Regression therefore corresponds to learning a 4096-weight-set *w*, that alters the importance of each feature so that the sum of the products estimates human similarity.

Several subsequent studies have extended this approach. Other forms of linear-transforms of the embedding matrix from images have been studied (Attarian et al., 2020). In computational linguistics, reweighting of word-embedding vectors improved prediction of human similarity judgments (Richie and Bhatia, 2020). Non-linear reweighting was also shown to be effective in modeling similarity judgments (Sanders and Nosofsky, 2020).

The effectiveness of reweighting speaks to strengths and limitations of vision-DNNs. It suggests that DNNs’ penultimate layer already forms a useful basis-set for predicting human similarity in that it captures the relevant features. Nonetheless, when emulating human similarity judgments, recalibration is required in order to assign features their correct levels of salience (see Richie and Bhatia, 2020, for similar argument in context of word embeddings).

### 1.2 Framework and Aims

The departure point of the current work is that the argument for reweighting may reflect an under-appreciation of the capacity of DNNs to predict human similarity judgments. Reweighting operationalizes the assumption, summarized above, that DNNs learn relevant features but assign them different levels of salience with respect to humans. A different possibility, which we probe here, is that DNNs do in fact acquire the relevant features at appropriate levels of salience. It is just that that in any particular evaluation context where human similarity-space is predicted, the contribution of relevant features is diluted by irrelevant ones. Specifically, taking the *entire* penultimate layer of a DNN as the relevant basis set effectively combines two representational sub-spaces: those relevant for human similarity judgments and those less relevant. While re-weighting can be considered as a way to de-mix these spaces, it comes with two costs. First, reweighting is applied via linear or non-linear transforms of node activation values in the DNN’s penultimate layer. This does not directly translate to a change in network architecture as no network weights are changed but instead a positive or negative multiplier is applied to the dot product of a feature’s values for two images. Explainability is further reduced by the fact that interpreting regression coefficients for reweighting is non-trivial even in the case of linear regression, and to our knowledge has not been attempted. Second, reweighting strongly impacts a network’s ability to classify, which increases the difficulty of understanding the relation between those features important for classification and those important for predicting similarity. To confirm that reweighting abolishes classification, we successfully reproduced the analyses and results of Peterson et al. (2018), and then evaluated VGG-19’s accuracy after the penultimate layer was reweighted. Top-1/top-5 accuracy dropped from {72.7; 91.0} to {9.4; 23.92} respectively.

The alternative we propose and evaluate is to identify a non-reweighted subset of features that is most informative for approximating a given set of similarity judgments obtained from human judgments or brain activity. The idea then is to use a pre-pruned DNN network to model human similarity. The intuition behind this idea borrows directly from neurobiology. Neuroscientists often quantify the 2OI between multivariate brain-activity and human similarity judgments, and this has strongly advanced knowledge of brain areas sensitive to particular categories. When doing so, one of the first steps is feature selection: selecting a limited set of brain-features (e.g., fMRI voxels or EEG sensors) that are expected to contain relevant information. For example, similarity judgments for scenes, objects, or faces would naturally be predicted using multivariate activation patterns sampled from different brain areas. That is, relatively limited brain areas will be selected, depending on the semantic categories studied, their breadth and depth. In some cases, an exhaustive “searchlight” search is conducted over the entire brain, within small volumetric parcels, to identify parcels where multivariate activity tracks similarity.

Conversely, it is *not* the case that the neurobiological feature set will consist of whole-brain activity, or even the entire visual/temporal cortex without further differentiation. Though vast areas of the brain are involved in even simple tasks (e.g., Gonzalez-Castillo et al., 2012), not all areas are equally relevant, and a preliminary selection of of brain areas is mandatory for obtaining neurobiologically-meaningful estimation. We put that the same holds when studying DNNs: it does not necessarily make sense to use all features (embeddings) for purposes of modeling a particular set of similarity judgments. This motivates Aim 1.

#### Aim 1

Learn pruned DNN configurations that improve out-of-sample prediction of human similarity judgments from a DNN’s penultimate layer, without activation reweighting. Pruning is implemented by feature selection that is supervised by human similarity structure. To the extent this aim was accomplished, we planned two derivative aims: Determine if pruned networks provide a better match to taxonomic structure (WordNet), and evaluate whether classification errors for pruned networks indicate biases (increased attention) towards the category optimized for similarity judgment.

#### Aim 2

Learn pruned DNN configurations that improve out-of-sample prediction of representational spaces manifest in human brain activity.

## 2 Study 1: Predicting human similarity judgments

### 2.1 Data set

The image set and similarity ratings data we use were curated by Peterson et al. (2018) and kindly provided to us by the authors. The image set consists of images from six categories, each with 120 images. The categories vary in perceptual and semantic heterogeneity. For consistency we refer to them using the labels introduced by Peterson et al.

1. ANIMALS: include photos of birds, reptiles and mammals of different types.
2. AUTOMOBILES: includes various transportation devices including sleds, rafts, trucks, trains, wheel barrels, planes, blimps, roller-skates, and horses.
3. FRUITS: fruits; mainly in context of original vegetation.
4. VEGETABLES: vegetables; mainly in context of original vegetation.
5. FURNITURE: mainly household furniture captured in indoor settings.
6. VARIOUS: a mix of images from the above categories but also including faces of people and outdoor scenery.

As evident some categories have greater taxonomic breadth than others, with VARIOUS being the broadest. Within each category, pairwise similarity ratings were obtained from humans for all combinations of 120 images; no similarity ratings were obtained for cross-category pairs, though the VARIOUS category can be treated as similarity mapping across superordinate-level categories.

### 2.2 Network pruning via supervised feature selection

We perform separate analyses for each of the 6 different categories. Each category contains 120 images. The human similarity-space is the upper-triangle of the 120×120 similarity-judgment matrix, and the DNN similarity space is the upper triangle of all pairwise cosine distances between images. For the DNN analysis, activation values were extracted from the penultimate layer of VGG-19, which contains 4096 nodes.

To improve prediction of human similarity ratings from DNN activity we implement a variant of a sequential feature selection (SFS) algorithm that is supervised by human similarity judgments. The process, implemented separately for each category, consisted of *i*) determining feature contribution, *ii*) selection-to-criterion, and *iii*) out of sample testing. The process was repeated for 5 folds in a cross-validation framework.

#### Determining contribution

In each cross-validation iteration, 20% (*n* = 24) of the images were designated as a test set and 80% (*n* = 96) as train set. *Baseline-2OI* was defined as the train-set’s 2OI between the DNN representation and human similarity judgments for those 96 images. We quantified each feature’s contribution to baseline-2OI by removing only that feature and recomputing train-set-2OI. The feature was then reinserted and the next removed till the process was repeated for all 4096 features. Consequently, ‘relevant’ features are those whose removal produces a 2OI value below baseline-2OI and ‘irrelevant’ features are those whose removal produces a 2OI value above baseline-2OI. This produces a rank order of each feature’s independent importance to baseline-2OI.

#### Selection-to-criterion

After ranking, we consecutively insert features, according to their importance rank, into a candidate feature set. Each time a feature is added to the set, we recompute 2OI against the train-set human similarity judgments. We add all features exhaustively and then we identify the set of features associated with the maximal value reached. The set thus identified constitutes the pruned network associated with a specific fold.

#### Out-of-sample generalization

once a pruned node-set is determined, we apply it to the left-out test set. The test-set images are inserted into the DNN, and coded as activation values for the retained nodes in the pruned layer. We then construct a pair-wise similarity matrix as described above and report pruned-net-2OI as *R*^2^. We evaluate this value in relation to the *test-set’s* baseline 20I which is the *R*^2^ produced when considering all 4096 nodes rather than the retained subset.

### 2.3 Improved prediction of human similarity

Pruning markedly improved prediction of human similarity judgments for out of sample image-sets as compared to the non-pruned Baseline. Results for each of the six categories are reported in Table 1. Out-of-sample prediction improved for all categories. The number of nodes required differed across categories, though there was no strong indication of a relationship between number of nodes maintained and the *R*^2^ achieved. For example, the ANIMALS and VARIOUS categories were the ones with most nodes maintained, but were associated with markedly different *R*^2^ values. For reference we also include values produced by other alignment methods when tested on the same cross-validation folds. It can be seen that pruning outperformed the other methods in 17 of the total 18 comparisons.

1. Baseline refers to the match between the DNN and human similarity space prior to any modification, averaged across the five out-of-sample data for the test folds.
2. PAG18 refers to application of ridge regression as implemented by Peterson et al. (2018), but applied to the five out-of sample folds used in our data.
3. Sim-DR is a reweighting approach developed by Jha et al. (2020) which optimizes a projection of DNN embeddings to a lower-dimensional space that matches human similarity judgments (we include values reported by the authors on the same dataset, as we have not implemented this learning model).
4. LASSO is our own variation of the reweighting implemented by PAG18 but which uses LASSO-regularized regression (Tibshirani, 1996) that is further constrained to only positive weights. As noted by others (Attarian et al., 2020) PAG18’s use of ridge regression produced negative weights which is psychologically inconsistent as it means there are features for which a larger product-term results in reduced similarity.
5. Pruned refers to the node-pruning method we introduce here.

**Table 1:** *R*^2^ for out-of-sample prediction of human similarity from pruned and original penultimate layer of VGG19 (baseline). For the pruned layer we also report the average number of nodes selected (± SD across folds).

|  | Animals | Automobiles | Fruits | Furniture | Various | Vegetables |
| --- | --- | --- | --- | --- | --- | --- |
| Baseline | 0.61 (0.07) | 0.51 (0.07) | 0.33 (0.08) | 0.29 (0.05) | 0.43 (0.10) | 0.32 (0.07) |
| PAG18 | 0.71 (0.09) | 0.50 (0.05) | 0.25 (0.15) | 0.34 (0.08) | 0.50 (0.13) | 0.27 (0.07) |
| LASSO | 0.64 (0.12) | 0.51 (0.08) | 0.38 (0.13) | 0.37 (0.11) | 0.47 (0.12) | 0.31 (0.08) |
| Sim-DR | 0.64 | **0.57** | 0.30 | 0.33 | 0.50 | 0.30 |
| Pruned | **0.75** (0.05) | 0.55 (0.08) | **0.39** (0.08) | **0.38** (0.07) | **0.56** (0.1) | **0.41** (0.05) |
| # nodes | 807 (63) | 647 (45) | 563 (76) | 557 (101) | 830 (44) | 605 (190) |

To obtain a better insight of the similarity spaces, we present the Multi Dimensional Scaling (MDS) plots of the non-pruned network and the network pruned for animals, on the 398 ANIMAL categories of ImageNet (Deng et al., 2012). The best exemplar image embeddings (as described in Section 2.4.1) for each of the 398 categories were used to construct the two dimensional MDS plot using sci-kit learn Python library with maximum iteration limit of 10,000 and converge tolerance of 1e-100. We chose the best fitting solution among four independent initializations. The images in the MDS plot are color coded to represent the six broad groupings of Animals within the WordNet tree. The results as presented in Figure 1 show that the pruned embeddings produce better-defined clusters, more tightly clustering fish, reptiles, amphibians and invertebrates (image details have been removed to comply with bioRxiv policies; full panels available at https://doi.org/10.6084/m9.figshare.14932725).

**Figure 1:**
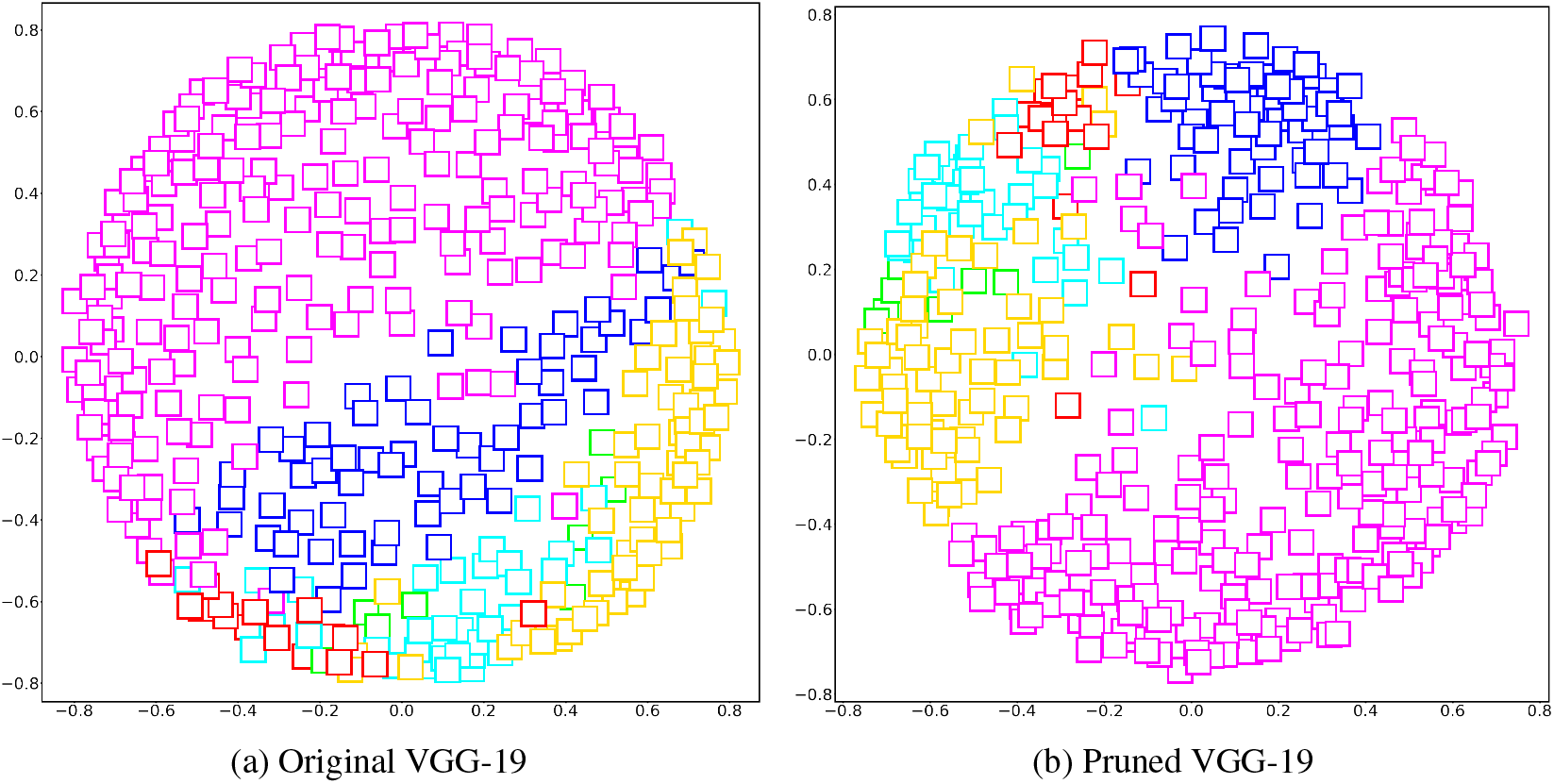
Multi Dimensional Scaling Plots of the embeddings corresponding to the 398 animal classes of ImageNet with Original VGG-19 and the same network pruned for Animals. Magenta-mammals, Yellow-invertebrates, Cyan-reptiles, Green-amphibian, Blue-bird, Red-fish.

### 2.4 Improved alignment with WordNet’s hierarchical taxonomy

To evaluate the impact of pruning on modeling taxonomic structure we focus on the ANIMALS category. The fact that pruning improves 2OI suggests that it maintains features that are important for separation between animal categories, but because 2OI reflects ranking of pair-wise similarities, it does not directly address heirarchical structure. To this end we compared the hierarchical structure latent in the DNN similarity space to that of WordNet (Fellbaum, 1998), which is a lexical database that also includes IS-A relations between animal types. We capitalize on the fact that ImageNet’s labels are derived from WordNet.

#### 2.4.1 WordNet/DNN Methods

To determine hierarchical information latent in the DNN similarity spaces we evaluated the relative fit between WordNet’s hierarchical structure and that of the pruned and non-pruned similarity spaces. From the DNN’s similarity spaces we derived hierarchical clustering analysis solutions from the pair-wise cosine distances. We used the scipy Python library, dendrogram with complete linkage function and the leaves within each cluster of the dendrogram were distance sorted in ascending order. To form the desired number of clusters from the dendrograms, we used *fcluster* from Scipy Python library with our criterion specified as maxclust.

To compute relative match with WordNet we used the Jaccard Index to assess to what extent cluster-members in the HCA solutions were also nearby-leafnodes in WordNet, as explained below. The input to the analysis were the 398 ANIMAL categories in ImageNet.

For the DNNs we computed HCA solutions from the embeddings in the penultimate layer associated with the best exemplar of each category. The exemplar was the image that produced the correct decision with highest confidence of all category members. From these embeddings we computed similarity matrices and HCA solutions with *N* = 6..12 clusters. To define the neighborhood-set of each category in the DNN’s HCA result, we extracted for each category member the set of all categories in the same HCA cluster.

To define the neighborhood-set of each category in WordNet we looked up the category in the WordNet graph, and extracted all leaf nodes subsumed by the category’s grandparent. This increased the granularity of WordNet as in many cases a category node had no siblings or very few ones, which made it non-feasible to use siblings as a neighborhood. This effective smoothing also usefully countered some ontological sub-divisions in WordNet that are not likely to have a counterpart in human similarity space (detailed below). The set-match was then computed per category by determining the Jaccard Index (JI) between the category’s DNN neighborhood-set and WordNet set (intersection of sets divided by union of sets). A grand mean was then computed over all categories.

WordNet contains multiple graph sections that increase in depth of IS-A links but without splitting (i.e, chains of parents that have only a single child; e.g., scorpaneoid → scorpaenid → lionfish). This is a knowledge-structure that does not appear to have a direct psychological analogue and can reduce the psychological validity of methods that use distances in WordNet as a proxy for conceptual distances (e.g., Huang et al., 2021). For the 398 categories we used, 100 were ones for which the target-node’s grandparent only subsumed a single leaf node (i.e., the target). For this reason we excluded these 100 categories from the analysis. We only analyzed categories with neighborhood-set sizes between 10 and 160, which resulted in using 245 categories of the total 398.

#### 2.4.2 WordNet/DNN results

In all cases, the pruned network’s hierarchical structure was a better match to WordNet than the non-pruned network. This was seen in that that Jaccard Index values were higher for the pruned network, and this held independent of the number of clusters set as a parameter (see Table 2).

**Table 2:**
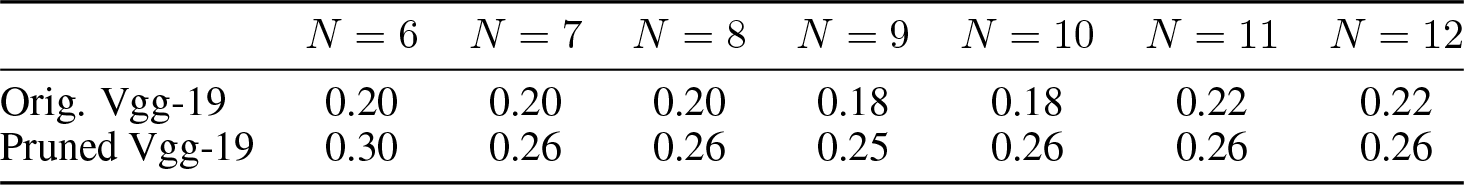
Jaccard-Index concordance between a category’s WordNet taxonomic neighborhood and its neighbors in DNN clusters. Higher values indicate greater agreement. Values shown for solutions across *N* = 6 : 12 DNN clusters. All comparisons statistically significant at *p* < .01 Bonferroni corrected for 6 comparisons.

To compute statistical significance we conducted an item-level paired analysis comparing the JI value for each category for the original and pruned cases. In all cases, JI values were higher for the pruned networks (Wilcoxon tests, Bonferroni corrected for multiple comparisons).

### 2.5 Pruning retains classification performance (without fine tuning)

The fully connected layers of DNNs contain substantial redundancy (e.g., Cheng et al., 2015), which suggests that pruned networks could retain adequate classification performance. We computed top1/top5 accuracy for pruned networks optimized for the six categories and report the mean values across folds (Figure 2). We find that top-5 accuracy never dropped below 79% and top-1 accuracy remained above 56%. Pruning provided much better performance than non-modified Ridge regression (Peterson et al., 2018). Pruning also outperformed our own implementation of Lasso-regression with a positive weight constraint, though with smaller margins.

**Figure 2:**
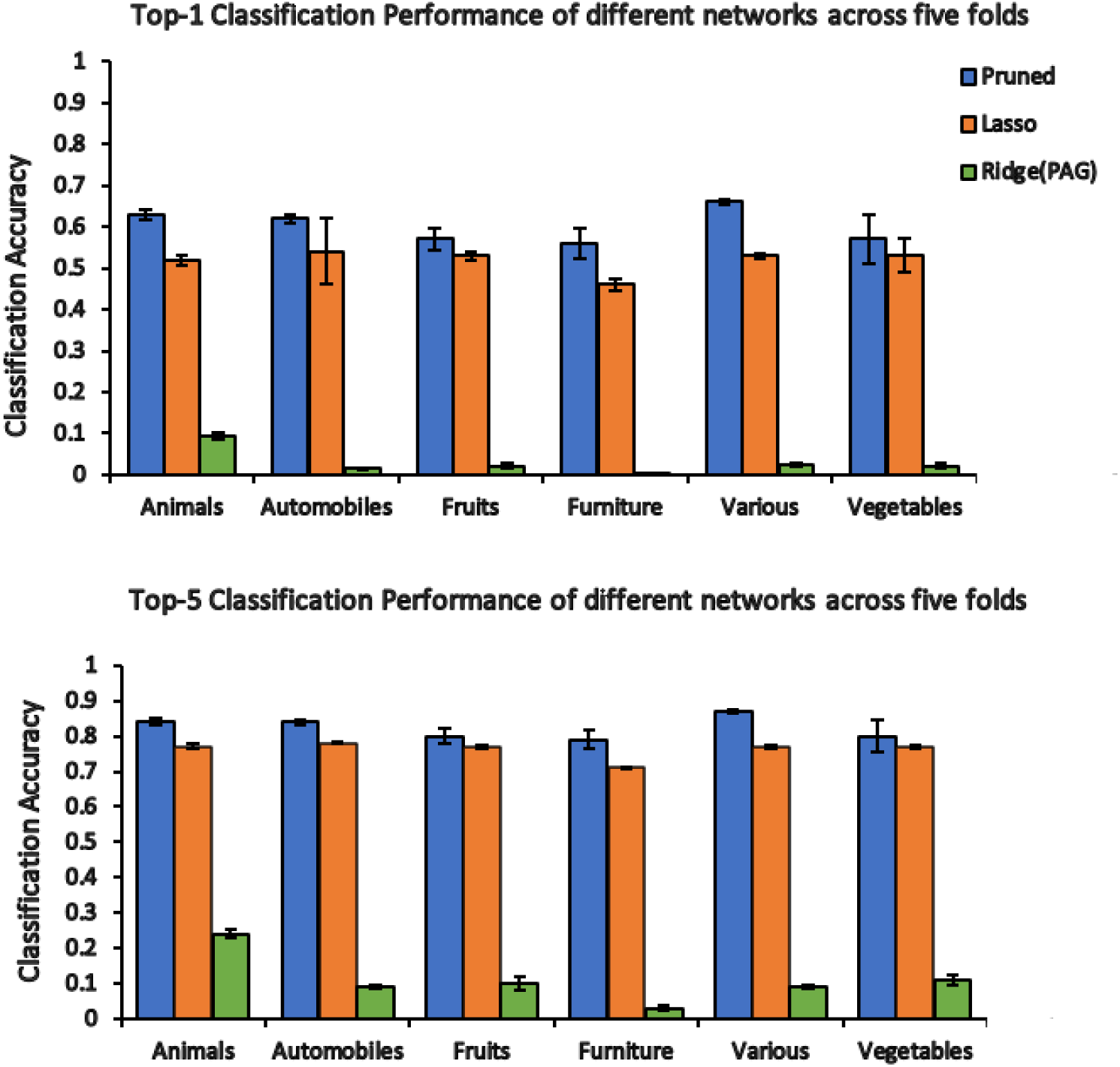
Classification performance of different networks for the different categories, averaged across the five folds. (a) Top-1 performance (b) Top-5 performance

### 2.6 Different instantiations of pruning produce similar activation patterns at penultimate layer

Because we established behavior of pruned networks across test-folds we could also determine to what extent different instantiations of the pruned layer, for a given category, produced similar activation patterns at the final (categorization) layer. Finding similar activations would suggest that different prunings (supervised by different sets of similarity judgments) still contained similar categorization-related information. For each category separately, we examined post-softmax final-layer patterns for pruned configurations produced in the 5 folds. Specifically, we used ImageNet’s 50K validation set. Per fold, we saved the 1000-valued vector of softmax outputs per image. Then, for each pair of two folds among the possible combinations of (5 choose 2), we computed the correlation between the softmax values for the same image across the two folds. We finally took the mean of those cross-fold correlations as a measure of correspondence at the final layer. We found substantial consistency in post-softmax activation vectors. For ANIMALS, Min/Max values for cross-fold correspondence were *R* = 0.92 − 0.96. For the other categories these values were, Vehicles: 0.90 − 0.95, Fruits: 0.85 − 0.94, Furniture: 0.86 − 0.96, Various: 0.94 − 0.96, Vegetables: 0.68 − 0.93.

### 2.7 Pruned networks shift error pattern

Pruning biases categorization errors towards the supervised category. We analyzed responses to ANIMAL images and focused on ‘substantial errors’, which we defined as trials where the true label was not among the top-5 post-softmax activations for a given input image. Using ImageNet’s 50K validation data set we identified images that constituted substantial errors for both the unmodified VGG-19 architecture and the pruned network (separately for each of the 5 folds/evaluations). When analyzing confusions within these substantial errors, we found that pruned networks more frequently classified images as animals. The magnitude of the bias, relative to the original VGG-19, appeared independent of whether the correct label belonged to the animal category or not. When the correct label was a sort of Animal, orig-vgg19 labeled the image as (the wrong) animal 69% of the time whereas the pruned-net in 91% of the time. When the correct label was not a sort of animal, the values were respectively 5% and 30%. The fact that the bias was independent of whether the correct label was an animal or not suggests the pruned network did not develop enhanced (useful) sensitivity to animals features, but expressed a linear (constant) bias to classify any image as an animal.

## 3 Study 2: Predicting neural similarity spaces

Pruning can be supervised by any *N × N* pair-wise similarity matrix. Here we evaluated if supervision of DNN pruning by neurobiological data increases the ability of DNNs to predict brain activity. Sequential Feature Selection was implemented identically to that reported above. The main difference was that instead of using human similarity judgments we used pairwise similarity values from multivariate neurobiological data collected using fMRI and Magnetoencephalography (MEG).

### 3.1 fMRI: Pruning improves prediction of neurobiological similarity space

#### 3.1.1 fMRI: Pruning methods

fMRI data and the image-materials were obtained from a public data set made available by King et al. (2019) who examined, in part, the second order isomorphism between DNN, human, and brain-derived similarity matrices. To maintain consistency with the neurobiology literature we use the term Representational Dissimilarity Matrix (RDM) which is a distance matrix computed by subtracting a pairwise similarity matrix from 1. The data set consisted of Human, DNN and Brain RDMs computed for two image-sets, each set consisting of 144 images from 48 categories. Different participants made similarity judgements for each set, making this dataset applicable for testing cross-participant generalization. Human RDMs were derived using an item-arrangement method that produces pairwise distances between images. Brain-derived RDMs were computed using typical multivariate analyses for different regions of interest (ROIs) associated with object and scene perception (see King et al. for details). This made it possible to use each ROI separately for supervising pruning. For the DNN, the authors report RDMs constructed from the VGG-S architecture but we re-implemented the analysis using the VGG-19 architecture. We note that RDMs produced by VGG-19 (penultimate layer) and VGG-S (final layer used by King et al.) were moderately correlated: for the two sets, values were 0.76, 0.78. Correlation between the DNN RDMs and behavioral RDM was almost identical for VGG-S and VGG-19 (in both cases, Pearson’s *R* ~ 0.6).

Before using the brain RDMs to supervise DNN pruning we evaluated whether we could replicate Study 1 in showing that pruning improves a DNN’s ability to predict human similarity judgements. For each set separately we used all pairwise similarity judgments between the 144 images to prune the DNN’s penultimate layer (*n* = 4096 nodes). Throughout our analysis, for each set, we used the mean behavioral RDM across all subjects within the set to supervise pruning. (We opted to prune using mean behavioral RDM reflecting all 144 images rather than mean behavioral RDM reflecting 48 categories in order to learn similarity relations within category.) Pruning, implemented via 5-fold cross-validation within each image set improved prediction of human RDMs for both Set1 and Set 2 (means across validation folds: Set1: *R*^2^ = 0.28 → 0.36; Set2: *R*^2^ = 0.22 → 0.30. In addition, because the two sets consisted of (different) images that belonged to the same categories, we could determine whether pruning the DNN based on the behavioral RDM of one image set improves 2OI for the other image set (a simple two-fold cross-validation). For both sets we found improved 2OI. Set1: *R*^2^ = 0.25 → 0.29 ; Set2: *R*^2^ = 0.22 → 0.25.

We evaluated eight brain areas for which data were provided: ventral temporal cortex (vTC), lateral occipitotemporal cortex (lOTC), fusiform face area (FFA), occipital face area (OFA), parahippocampal place area (PPA), occipital place area (OPA), and ventral and dorsal early visual cortex (vEVC and dEVC).

We implemented four tests of pruning. The first pruned the DNNs based on behavioral RDMs and evaluated the impact on predicting brain RDMs. In contrast, analyses 2, 3 and 4 used the brain RDMs themselves to supervise pruning of the DNN. To maintain compatibility with prior literature (e.g, King et al., 2019), these latter analyses also considered both the penultimate and final (*n* = 1000 nodes) layers of the network.

In the first analysis, we pruned DNN nodes using the behavioral RDMs (one RDM per image set) to evaluate if this will improve the 2OI between DNN and brain RDMs. In this case, for each image set a single pruned network configuration was determined by the best fit between a pruned DNN RDM and behavioral RDM. This pruned configuration was used to derive (pruned) DNN RDMs that were compared with the brain region RDMs.

In the second analysis, within each image set separately, we pruned either VGG-19’s penultimate layer or VGG-19’s final layer to determine if pruning improves prediction of brain RDMs for out of sample images (within-set cross validation). For each brain ROI, we employed 5-fold cross-validation where we supervised pruning based on group-level RDM (average of all single-participant RDMs within each set separately). In each fold, 80% (n= 115) of the 144 images composed the train set, and the remaining composed the validation set. There was no overlap between the images used for training and validation. To evaluate the impact of pruning, 2OI was computed for the validation subset in each fold, both for the pruned and non-pruned embeddings.

In the third analysis, we used the 2-fold approach that we had applied for the behavioral data where we pruned DNNs for one image set, and then evaluated the pruning on the other image set (cross-set prediction).

In the fourth analysis, we implemented pruning outside a cross-validation context, where the complete brain RDM (from all 144 images) supervised pruning of a complete DNN RDM constructed from the same images. While this can be considered as over-fitting, it can also be seen as an upper-bound indicator of the ‘signal’ contained in a given DNN layer with respect to its ability to predict a brain RDM, making it an important quantity.

#### 3.1.2 fMRI: Pruning results

Our first analysis showed that pruning a DNN based on a behavioral RDM improved the isomorsphism between the DNN RDM and RDMs of the 8 regions of interest. Table 3 presents the 2OI *R* values for the raw and pruned DNNs. A paired t-test applied to the Fisher-Z transformed *R* values (considering the 16 comparisons; 8 regions per two sets) confirmed these impressions statistically, *t*(15) = 2.88, *p* = 0.005 (one tailed for *pruned > raw* directional test). This shows that even a very general pruning based on human similarity judgements can improve 2OI with brain ROIs. As indicated above, the remaining analyses used each brain region’s RDM separately to supervise DNN pruning.

**Table 3:** Pruning constrained by human similarity-judgments improves 2OI *R* between DNN RDMs and Brain-region RDMs.

|  | vTC | IOTC | FFA | OFA | PPA | OPA | vEVC | dEVC |
| --- | --- | --- | --- | --- | --- | --- | --- | --- |
| Set 1 | 0.130 | 0.099 | -0.021 | 0.078 | 0.184 | 0.098 | 0.022 | 0.041 |
| Set 1 pruned | **0.152** | 0.097 | **-0.018** | **0.089** | **0.207** | 0.089 | **0.054** | **0.060** |
| Set 2 | 0.150 | 0.073 | 0.009 | 0.066 | 0.187 | 0.139 | 0.023 | 0.017 |
| Set 2 pruned | 0.146 | **0.079** | **0.027** | **0.071** | 0.187 | 0.134 | **0.033** | **0.019** |

Figure 3 shows results for within-set cross validation. It shows that learning DNN pruning from brain RDMs was highly effective in increasing the ability to predict brain RDMs for out of sample images. In evaluating the brain ROIs we found improved prediction for 31 of the 32 analyses (pruning evaluated for 8 regions, for two image sets, for two layers). To evaluate the data statistically, we analyzed results for each layer separately. For the penultimate layer We contrasted the 16 raw correlation with the 16 values from the pruned network (after Fisher-Z transform). The mean values were differed markedly (*M*_*pruned*_ = 0.127 vs. *M*_*raw*_ = 0.0798) and the difference was statistically robust accompanied by a large effect size, *t*(15) = 7.02, *p* < .001, *d* = 1.75. A similar result held for the final layer, *M*_*pruned*_ = 0.158 vs. *M*_*raw*_ = 0.0873), *t*(15) = 6.06, *p* < .001, *d* = 1.51.

**Figure 3:**
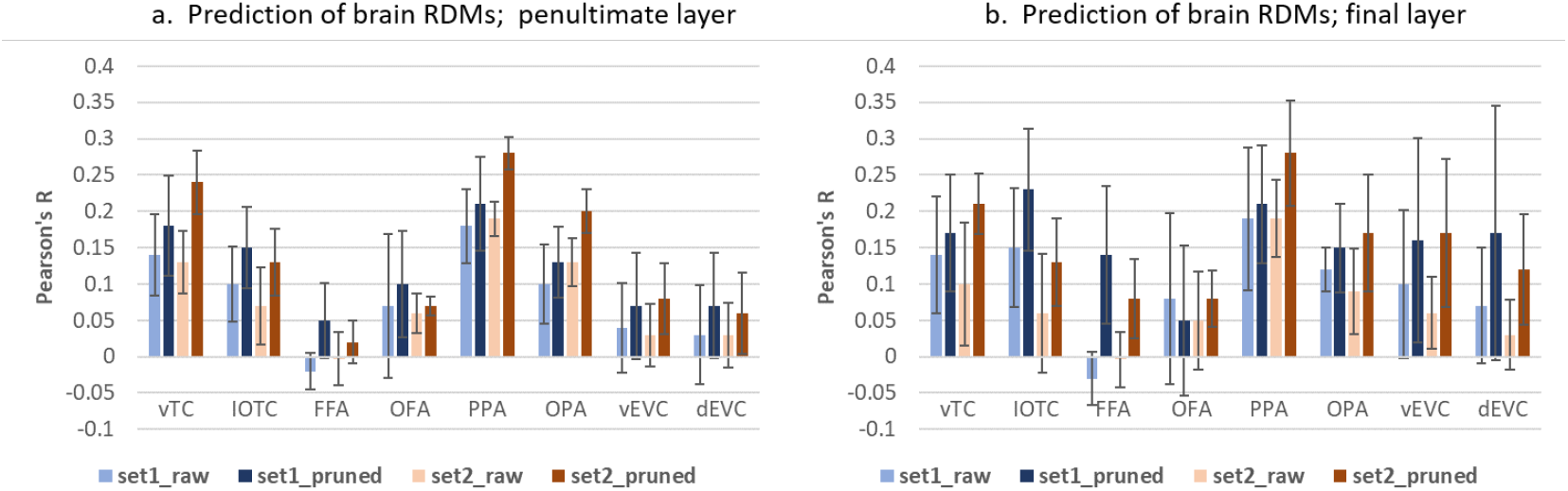
Learning pruning within each image set for separate cortical ROIs. Pruning was tested using 5-fold cross validation, separately for image set 1 and image set 2. (a) Prediction from embeddings from VGG-19 penultimate layer (b) Predictions from embeddings from VGG-19 final (1000-node) layer

As in King et al. (2019), we found relatively weak results for the FFA when using the complete embeddings (for both the penultimate and final layer), but supervised pruning improved the prediction of this region’s RDM, particularly when applied to VGG-19’s final layer (see Figure 3b). Qualitatively, for vTC pruning of the penultimate layer was more effective than pruning the final layer, but for FFA, vEVC and dEVC a converse pattern held suggesting these regions benefit from pruning information coded at the category level of the final layer. As we discuss below, this could occur whenever a distribution of values in the final layer constitutes an effective lower-dimensional space.

Figure 4 reports results of the cross-set RDM prediction, which also generalizes across any between-participant variance. Here too pruning improved prediction of brain RDMs almost without exception (30/32 of cases examined). And as in the within-set analysis, pruning improved the ability to predict FFA RDMs, and more strongly so when using the final layer. For the penultimate layer the mean values differed (*M*_*pruned*_ = 0.130 vs. *M*_*raw*_ = 0.081), *t*(15) = 9.37, *p* < .001, *d* = 2.34. The same held for the penultimate layer, (*M*_*pruned*_ = 0.137 vs. *M*_*raw*_ = 0.088), *t*(15) = 4.30, *p* < .001, *d* = 1.07.

**Figure 4:**
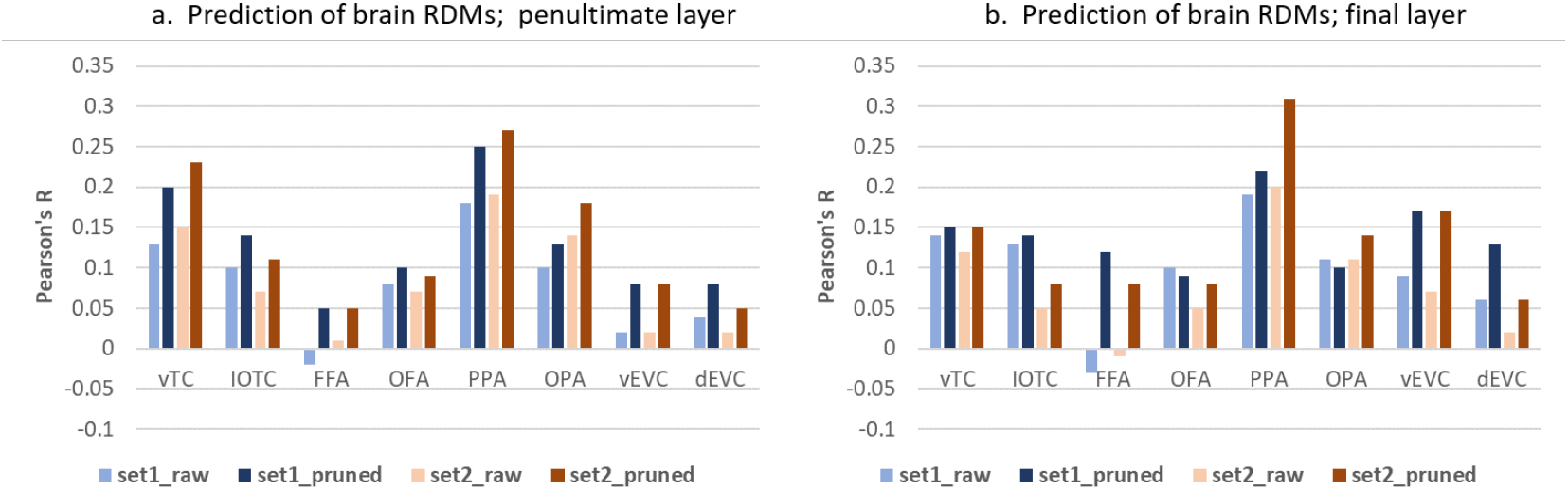
Learning pruning across image sets. Pruning was learned for DNN embeddings for one image set and applied to the other image set. (a) Prediction from embeddings from VGG-19 penultimate layer (b) Predictions from embeddings from VGG-19 final (1000-node) layer

Finally, we evaluated the impact of directly pruning DNN embeddings based on brain RDMs outside a validation context (Figure 5). This shows to what extent pruning improves the match between DNN and brain RDMs when applied to embeddings produced from the same image set. As seen in the Figure, pruning improved isomorphism between DNN and brain-region RDMs across the board, often by substantial multipliers.

**Figure 5:**
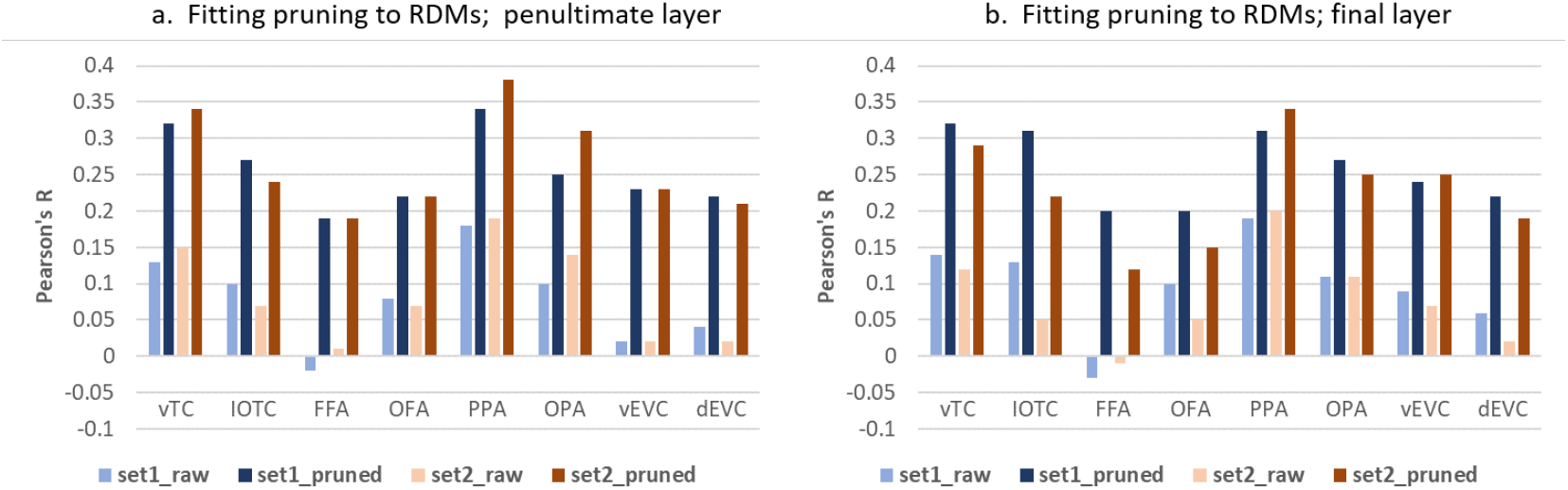
Direct pruning of DNN RDMs from brain-region RDMs without generalization. (a) Fitting VGG-19 penultimate layer (b) Fitting VGG-19 final (1000-node) layer

Importantly, this last analysis also allowed us to determine how many features were retained in optimizing the fit between RDMs. as shown in Table 4, when applied to final layer of VGG-19 (*n* = 1000 nodes), pruning generally selected a low number of nodes. What immediately stands out in the Table is that pruning DNN embeddings from FFA RDMs was optimized by selecting as few as ten (or less) nodes, for both image sets. In contrast, PPA required a substantially larger number of nodes and was associated with the largest number of retained nodes for both image sets. There was a moderate agreement between the number of nodes retained across sets (*n* = 8 regions, Pearson’s *R* = 0.52) indicating that the number of nodes retained through pruning, per brain ROI, is systematically linked to the information coded for in that brain area.

**Table 4:** Number of nodes retained from VGG-19 final layer for pruning based on different ROIs.

|  | vTC | IOTC | FFA | OFA | PPA | OPA | vEVC | dEVC |
| --- | --- | --- | --- | --- | --- | --- | --- | --- |
| Set 1 | 27 | 29 | 10 | 60 | 75 | 40 | 41 | 71 |
| Set 2 | 40 | 79 | 7 | 128 | 138 | 84 | 12 | 18 |

It is important to keep in mind that pruning embeddings from the final layer does not mean that a given brain region is necessarily sensitive to defining features of the retained nodes. It only means that the multivariate activity pattern in those nodes across images, as expressed in a DNN RDM, tracks the brain-region’s RDM. This multivariate activity pattern would reflect any meaningful covariance between the penultimate layer and final layer. For example, for Set 2, the brain RDM for FFA was optimized by selecting only 7 of the 1000 nodes, and these 7 nodes had the following labels: *geyser, volcano, killer-whale, steel-arch-bridge, steam-locomotive, electric-locomotive, strainer*. This just means those 7 nodes constitute a useful lower-dimensional space for tracking regional-FFA response, and the reason for this needs to be explored using methods suitable for studying lower dimensional spaces in DNNs.

### 3.2 MEG: applying pruning to different processing stages

#### 3.2.1 Methods: learning pruning from MEG data

Participants in the MEG study (*n* = 20) observed images drawn from 160 basic-level categories, among them multiple types of animals which are our focus here. Acquisition rate was 200Hz and data of interest were those collected during 1sec beginning 200ms prior to stimulus presentation and ending 800ms afterwards. This produced 200 observations at 5Hz, for each of 360 MEG sensors. The MEG-derived similarity for a pair of images {*k, j*} was computed at each time point *t*_1_ to *t*_200_. Similarity was defined as the correlation between the MEG activation vectors of images *k* and *j* over all 360 sensors. This is the regular procedure using MEG (e.g., Giari et al., 2020) as each MEG sensor reflects contributions of multiple activation sources in the brain, so that activity is not strongly localized topographically.

Rotating over all image combinations, this produces a representational dissimilarity matrix (RDM) at each time point *t* which is the analogue to that derived from human similarity judgments or fMRI-ROI RDMs above. We evaluated the fit of this RDM with that produced from the original VGG-19 RDM, and that produced from pruned RDMs via sequential feature selection.

Based on prior literature, we computed the fit for 2 time windows, per person: One window between 175 and 185ms post stimulus, and the other between 275 and 285ms post stimulus. Within each window we averaged the the MEG cross-similarity matrix of all time points. Five-fold cross-validation was implemented per time point per subject. A time-window of interest approach was adopted due to computational processing constraints (the process requires around 45 minutes per time point per participant on a 16-core machine). The computed R values were Fisher-Z transformed and analyzed using a paired T-test (directional, *pruned > non* − *pruned*).

#### 3.2.2 MEG Results

For the 175-185ms post-stimulus window we did not find a statistically significant difference between *R* values for the pruned and non-pruned DNNs when applied to the validation set. Means of Pearson’s R value were similar and low (Pearson’s R; M = 0.004 vs. 0.002, *t*(19) = 0.82, *p* = 0.21).

For the 275-285ms time window, pruning improved 2OI, (Pearson’s R; M = 0.01 vs. 0.0045, *t*(19) = 1.99, *p* = 0.03, *d* = 0.46). We note that these mean values, while low, are completely consistent with prior literature on 2OI between RDMs produced from DNNs and MEG data (e.g., Giari et al., 2020) and the effect size at the group level (d = 0.46) suggested a substantial impact of pruning even in a domain with essentially low 2OI.

To conclude, we find that MEG supervised pruning increased the predictive power of DNNs with respect to MEG data collected around 280ms rather than earlier (180ms) processing periods, although the effects are less robust compared to measures with inherently higher signal to noise.

## 4 Limitations

Given that behavior-supervised pruning has not been implemented to date, the current effort exhibits limitations. From an implementation perspective, we implemented pruning as an additional machine learning step rather than integrating it into the DNN training itself. Future work could integrate supervised pruning with the classification training as in Piggyback (Mallya et al., 2018). This is particularly important if one intends to target pruning of convolutional layers below the fully connected ones. Those layers have a much higher dimensionality (hundreds of thousands of nodes), which can make sequential feature selection non viable. It would also be important to evaluate the efficacy of pruning for other DNN architectures. Empirically, supervision by fMRI RDMs was robust, but the supervision by MEG RDMs was weaker, which likely reflects the low signal-to-noise in MEG data. Typically-reported 2OI *R*^2^ values between DNN and MEG data in the literature are modest (Pearson’s R < 0.1). We had hoped to substantially improve on these values via pruning but ultimately find it is likely that MEG data are a low signal-to-noise domain that constitutes a weak supervisor for learning.

## 5 Discussion

We find that supervised-pruning of DNN nodes improves out of sample prediction of human and neurobiological similarity spaces. When direct comparisons were possible, pruning outperformed previous reweighting-based methods for modifying network activation (Table 1). In addition, hierarchical clustering applied to pruned-embeddings proved a better fit to WordNet’s hierarchy, and MDS spaces produced from pruned embeddings reflected a better clustering. For neurobiological data, pruning substantially improved the ability to predict brain-region RDMs for out of sample images, and was able to extract signal from low signal-to-noise MEG data. These practical advantages however are secondary to the theoretical implications of pruning’s success. These indicate that DNNs already capture features relevant to human similarity spaces at an adequate level of salience, and for this reason, node activations do not need to be reweighted. One just needs to filter out those features/nodes that are less relevant to modeling similarity of the domain at hand. Supervised pruning therefore improves the sensitivity of quantifying isomorphism between DNNs and humans, and opens the door to new studies of semantic knowledge in artificial and biological systems.

## Acknowledgments and Disclosure of Funding

Scott Faiirhall is funded by the European Research Council (ERC) Starting Grant, ‘CRASK’, awarded under the European Union’s Horizon 2020 research and innovation program (Grant Agreement No. 640594).

